# Top-down contextual requirement to suppress self-movement influences grasp-specific modulation of corticospinal excitability during action observation

**DOI:** 10.1101/2020.06.25.171629

**Authors:** Steven J. Jerjian, Marco Davare, Alexander Kraskov

**Affiliations:** Department of Clinical and Movement Neurosciences, UCL Queen Square Institute of Neurology, London, United Kingdom; Department of Clinical Sciences and Centre for Cognitive Neuroscience, Brunel University London, United Kingdom; Biosciences Institute, Faculty of Medical Sciences, Newcastle University, NE2 4HH, United Kingdom; Zanvyl Krieger Mind/Brain Institute, Johns Hopkins University, Baltimore, Maryland, United States

**Author notes:** **Authors Contributions:** S.J.J. performed experiments, analysed data, and wrote the paper. S.J.J., M.D. and A.K. designed research, discussed results and conclusions, and edited the paper.

## Abstract

Action observation modulates corticospinal excitability (CSE) measured via transcranial magnetic stimulation (TMS) in humans, which presumably exposes the effect of mirror neuron activation on corticospinal pathways. These responses can consist of both facilitation and suppression, and the balance of these two may restrict the outflow of activity into movement. Evidence also suggests that task context can considerably influence CSE changes during action observation.

Here, we assessed whether embedding action observation within a Go-NoGo paradigm, emphasizing movement withholding on observation and NoGo trials, influenced CSE modulation. Fourteen healthy subjects received single pulse TMS over left primary motor cortex (M1) during a baseline period, grasp observation onset, or after a NoGo cue, while performing, observing, or withholding two distinct reach-to-grasp actions. We assessed modulation of motor evoked potentials (MEPs) in three intrinsic hand muscles, which were recruited in a grasp-specific manner during action execution. CSE modulation was limited, and predominantly suppressive in nature during grasp observation. Seven subjects performed the same task without the NoGo condition (“Go-only” block) immediately before the “Go&NoGo” block. We found evidence for grasp-specific modulation of CSE, which matched the recruitment pattern of the muscles during action execution. Within these subjects, modulation was attenuated when the NoGo condition was introduced, but was still distinct from modulation in the first group.

These results suggest that bottom-up grasp-specific modulation of MEPs during action observation is attenuated by the top-down contextual requirement to suppress self-movement, and facilitation and suppression effects are determined by the balance between these two processes.

**SIGNIFICANCE STATEMENT:** Action observation can activate specific pathways in the motor system of the observer, which are also used to perform the same action. This motor resonance, measured via changes in corticospinal excitability using transcranial magnetic stimulation, is susceptible to task context. In this study, we show that observing grasping actions results in grasp-specific changes in excitability, consistent with mirror neuron activation, but this effect is masked when observation is interleaved within a Go-NoGo paradigm, which emphasises suppression of one’s own movement. Top-down task requirements to withhold movement within and across trials, which are present in most action observation studies, likely influence the extent of motor resonance, urging caution in the design and interpretation of results in TMS action observation experiments.

## INTRODUCTION

The observation of another’s actions induces activity within motor pathways, even in the absence of self-movement. This activity is driven by mirror neurons, which modulate their firing rate during both execution and observation. Mirror neurons were first identified in the macaque ventral premotor area F5^1^, and have since been shown to exist in a range of frontal and parietal areas^2–5^. These neurons have been hypothesized to mediate a mapping of observed actions directly onto the motor system of the observer. Mirror activity even extends to pyramidal tract neurons^3,6^, which project directly to the spinal cord, and in humans, transcranial magnetic stimulation has demonstrated modulation of corticospinal excitability (CSE) during action observation^7–9^. Importantly, these findings provide evidence for both facilitation and suppression of activity in corticospinal pathways during action observation, and the balance of the two could be critical for preventing the outflow of cortical mirror activity into overt movement^3,10^.

A number of TMS studies have shown that the direction and magnitude of CSE modulation during the grasping phase of action observation depends on parameters, such as the force^11^ and kinematics^8,12^ of observed object-oriented actions. Furthermore, CSE modulation is affected by task instructions or requirements^13–15^, as well as the availability and quality of contextual visual information, either regarding the object to be grasped ^16,17^, or the acting agent ^18,19^.

One contextual requirement, present in almost all action observation studies, is the requirement to suppress self-movement while observing actions. In TMS studies, subjects are generally required to be at rest since voluntary EMG activity inflates MEP amplitudes. Similarly, since macaque mirror neurons respond to both execution and observation, voluntary EMG during action observation would confound accurate identification of mirror neurons^6^. On the other hand, there has been some speculation that this somewhat artificial requirement to stay relaxed, at least in human studies, may mask or otherwise influence CSE changes during action observation^13,20^, and that, importantly, the level of instruction or emphasis on this requirement may contribute to some of the discrepancies in results between studies in terms of facilitation or suppression of CSE.

In this study, we investigated whether further emphasis on actively suppressing one’s own actions, via the presence of NoGo trials embedded within action observation paradigm, could influence CSE modulation during action observation. The observed grasps took place within peri-personal space, which appear to be treated differently from actions in extra-personal space within the mirror neuron system^21,22^, and may encourage suppression of self-action. We hypothesized that the emphasis of the Go-NoGo task in particular on suppression of self-movement might influence CSE modulation during action observation. One group of subjects therefore performed a “Go-only” task involving only execution and observation, prior to performing the “Go-NoGo” task, to assess whether grasp-specific modulation during the “Go-only” task might be attenuated by the addition of the NoGo condition due to a shift in the task context.

## MATERIALS & METHODS

### Participants

14 right-handed subjects (21-41 years old, mean ± standard deviation: 29.4 ± 5.1, 8 female) participated in this study. Subjects were tested in two groups of seven (Group 1: 2 male, 5 female, age 28.3 ± 4.4, Group 2: 4 male, 3 female, age 30.4 ± 5.9). All subjects passed a screening test for contraindications to TMS and were neurologically healthy, with normal or corrected-to-normal vision. Subjects gave informed consent and were financially compensated for their participation. Experimental procedures were approved by the Local Ethics Committee.

### Experimental set-up

Subjects sat opposite an experimenter, rested their hands on two homepads, and viewed two objects through a controllable LCD screen (14×10cm), which remained opaque in between trials (Fig 1a). A marble object afforded precision grip (PG) via opposition of thumb and forefinger, and a sphere object afforded whole hand grasp (WHG) with all fingers encompassing the object (Fig 1a, inset).

**Figure 1.**
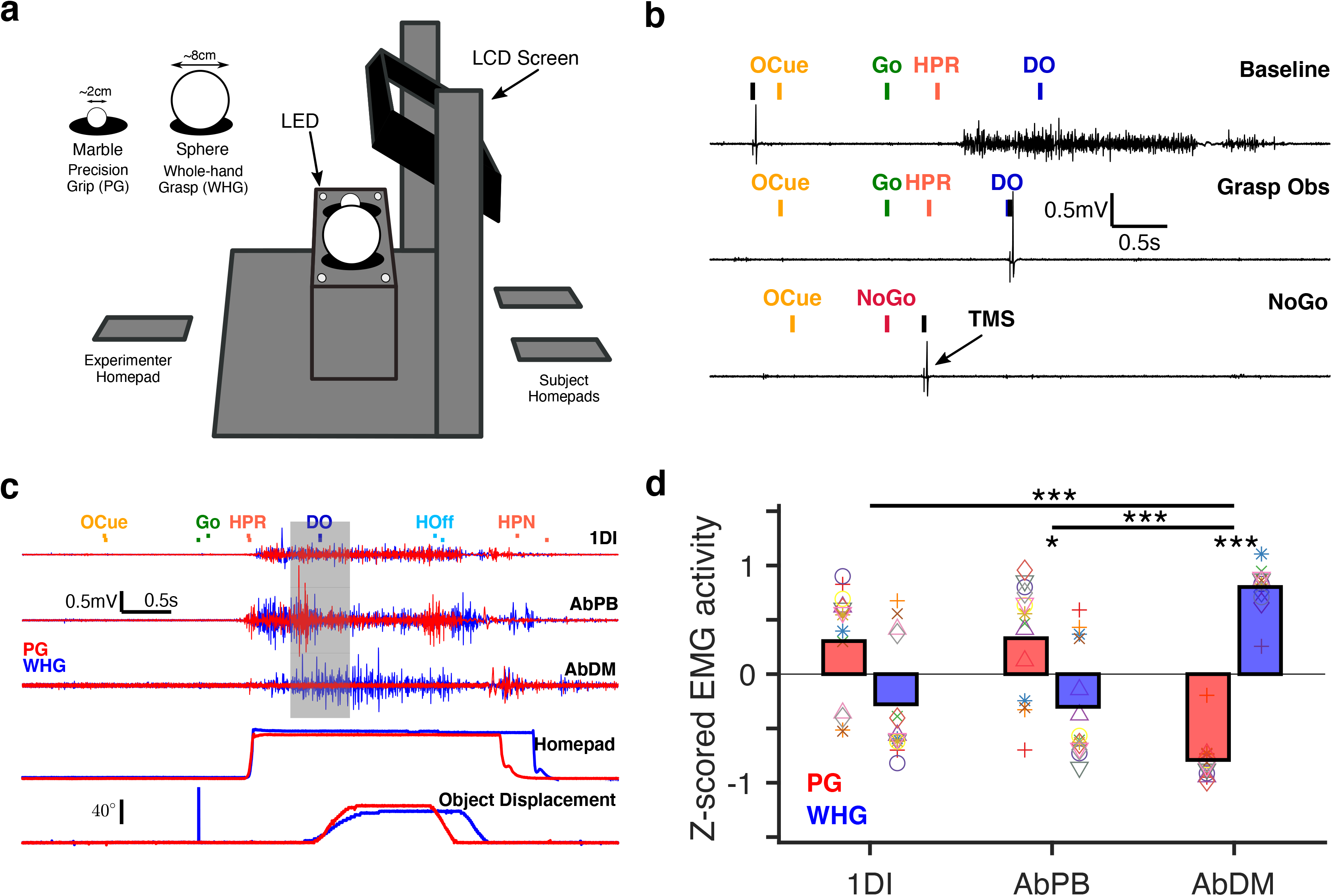
**(a)** Schematic of experimental box used for task (not to scale). Inset shows marble and sphere objects used for precision grip (PG) and whole-hand grasp (WHG), respectively. **(b)** Example raw EMG activity in 1DI muscle during one precision grip trial of each TMS condition from a single subject. TMS times for each condition are denoted by black vertical lines, and “TMS” label in bottom trace indicates TMS artefact and subsequent MEP. Coloured lines denote salient task event times indicated by labels (OCue – object cue, Go/NoGo – imperative cue, HPR – homepad release, DO – displacement onset). Trace labels (Baseline, Grasp Observation, NoGo) refer to the TMS trial type. Baseline TMS (delivered at the time the LCD screen initially became transparent at the start of the trial) could be delivered on execution or observation trials, which were indistinguishable at this timepoint (example shown for execution trial). Grasp Observation TMS was delivered concurrent with experimenter DO, and NoGo TMS was delivered 350ms after the NoGo Cue, intended to be close to the time of experimenter movement onset (HPR) during observation trials. **(c)** Raw EMG activity in each muscle, and homepad and object displacement signals during a single execution trial for each grasp (red – precision grip (PG), blue – whole-hand grasp (WHG)). Vertical pulse at Go cue on WHG object displacement channel corresponds to TTL pulse used for offline alignment of digital events and analog data. Vertical tick marks denote key event times for PG (first row) and WHG (bottom row). Grey shaded area denotes interval (±300ms around DO) used for average EMG shown in (d). **(d)** Z-scored EMG activity for different grasps and muscles, all 14 subjects. Bars show mean, individual subjects shown by symbols. * p < 0.05, *** p < 0.0001 in post-hoc comparisons.

Trials began when the subject depressed both homepads and the experimenter depressed the homepad on their side. The LCD screen became transparent, and the object area was illuminated. After a variable delay (uniformly sampled from 0.25-0.45s), two amber LEDs illuminated on one or other side to indicate the target object for the upcoming trial (**o**bject **cue**; OCue). After a further variable delay (uniformly sampled from 0.8-1.2s), a single green or red LED indicated the trial type (Go or NoGo, respectively). A green LED on the subject side indicated an execution trial. The subject released their right homepad (**h**ome**p**ad **r**elease; HPR), and used their right hand to make a reach-to-grasp movement towards the target object. After grasping the object using the required grasp (**d**isplacement **o**nset; DO), they rotated the object into a window ( > 30degrees), and held the grasp for 1 second. A constant frequency tone indicated that the subject had entered the hold window, and a second, higher frequency tone signalled successful completion of the 1 second hold. The subject then released the grasp (**h**old **off**; HOff) and returned their right hand to the homepad (**h**ome**p**ad retur**n**; HPN). Recorded muscle activity, homepad and object displacement signals relative to task events from an example trial are shown in Figure 1c. A green LED on the experimenter side, indicating an observation trial, required the subject to remain still, and simply attend as the experimenter performed the same reach-to-grasp action with their right hand. A red LED on the subject side indicated a NoGo trial, in which the subject (and experimenter) simply refrained from movement. After 1 second, a tone indicated successful completion of the NoGo trial. A short intertrial interval (1-2s) occurred between each trial, and all trial types were presented in a pseudo-randomised order. Execution trials were relatively infrequent to reduce the length of the session but were still included to assess muscle recruitment during action execution, and to encourage subjects to remained attentive throughout the task in case they were required to move. In blocks with the NoGo condition (“Go & NoGo” experiment; see “TMS and experimental procedure”), execution, NoGo, and observation trials were interleaved in a 1:2:3 ratio. In blocks without the NoGo condition (“Go only” experiment), execution and observation trials occurred in a 1:3 ratio.

### TMS and experimental procedure

Single-pulse TMS was delivered via a Magstim 200 stimulator attached to a standard figure-of-eight coil (70mm diameter). The coil was held tangentially to the scalp with the handle pointing backwards at a 45°angle to the midline, and the motor hotspot was identified as the position over left M1 eliciting the most consistent MEPs in the three muscles. Stimulator output was set to obtain MEPs averaging approximately 1mV in the resting 1DI muscle.

Single-pulse TMS was delivered on individual trials (Fig 1b.) at one of three possible time points (1) LCDon (within-task baseline), on either execution or observation trials, (2) Experimenter DO (Grasp Observation) on observation trials only (3) 350ms after the red LED on NoGo trials (NoGo, approximately just before experimenter HPR on observation trials). TMS timing was determined by triggers sent from the experimental box software (e.g. the Grasp Observation condition trigger occurred when the experimenter contacted the object), ensuring that the timing of stimulation was consistently at the intended time for a given condition. To ensure subjects could not anticipate stimulation, no TMS was delivered on 20% of trials. These trials were interleaved in a pseudo-random manner, and each trial therefore contained either zero or one TMS pulse. To maintain attention and limit fatigue, subjects completed the experiment in two shorter blocks, with approximately 10 MEPs per condition per block, resulting in a total of 20 MEPs per condition. To assess whether the presence of the NoGo condition may have influenced the implicit goal of the task and therefore also the modulation of CSE, seven subjects tested later completed an additional two blocks of the same task, which did not include a NoGo condition, but was identical in all other aspects. These “Go-only” blocks were always completed before the “Go & NoGo” blocks, and subjects were not introduced to the NoGo condition until these first blocks were completed, in order to keep them immune from any effect of explicit NoGo trials. In addition, prior to each block, a set of 20–30 baseline MEPs were recorded, with the subjects at rest (pre-task baseline), and single-pulse TMS delivered every 5±1s. Before starting the experiment, subjects were briefly familiarized with the task via verbal instructions and a short practice session, to check EMG signal quality and task performance.

### Recording

Electromyographic (EMG) recordings were made from 3 right hand muscles (Fig. 1c) (first dorsal interosseous (1DI), abductor pollicis brevis (AbPB), and abductor digiti minimi (AbDM), using surface electrodes mounted in a belly-tendon montage, with a ground electrode fixed to the bony prominence of the wrist. EMG signals were amplified (×1000), high pass filtered (3Hz) (Neurolog EMG amplifier NL824, and Neurolog isolator NL820, Digitimer Ltd, UK), and digitised at 5kHz. Concurrently, we recorded the timing of all task events and TMS triggers, and analog signals of homepad pressure and object displacement (5kHz). All data were stored on a laboratory computer for offline analysis.

### Data Analyses

#### Task behavior

Average reaction times for each subject and experimenter were computed as the median difference across trials between the green LED go cue and movement onset, when pressure on the homepad was released (HPR), for each grasp separately. Movement times were computed as the median difference between HPR (movement onset) and DO (object displacement onset).

### Statistical analysis of EMG activity during action execution

To quantify the extent to which subjects used each of the three muscles to perform the two grasps (Fig. 1c, red and blue traces correspond to PG and WHG, respectively), EMG signals were high-pass filtered (0.5Hz, 2nd order Butterworth), rectified, and low-pass filtered (30Hz, 2nd order Butterworth). EMG activity in the −300 to +300ms period around DO was then averaged to obtain a single value for each execution trial. This interval was chosen to capture the dynamic period in which hand shaping and grasp occurred. For comparison across subjects with different levels of EMG, values within each muscle were z-scored, and averaged for each object. The resulting values were statistically compared via 2-way repeated-measures ANOVA (rmANOVA), with within-subject factors of object (PG, WHG) and muscle (1DI, AbPB, AbDM), followed by post-hoc tests with correction for multiple comparisons (Tukey’s honestly significant difference (HSD)).

### Statistical analysis of MEPs

For each subject independently, MEPs were discarded from analysis if; 1) the mean rectified EMG activity in the 100ms preceding the TMS stimulus exceeded 2 standard deviations of the mean pre-stimulus EMG, 2) the peak-to-peak MEP amplitude exceeded 3 standard deviations of the mean MEP amplitude across all conditions in a given block, 3) the MEP amplitude did not exceed 0.05mV. Across all subjects, 5.8±0.3% of MEPs (mean±SEM across subjects) were discarded in the “Go & NoGo” experiment, and 5.0±0.4% of MEPs in the “Go only” experiment. The peak-to-peak amplitude of remaining MEPs for each subject were averaged across all trials for each condition, to obtain one value per TMS condition, grasp, and muscle for each subject.

To test for non-specific facilitation relative to the pre-task baseline, we compared the raw amplitude MEPs in the pre-task baseline period to those during the within-task baseline and grasping observation, using a two-way repeated-measures ANOVA (rmANOVA) with factors MUSCLE (1DI, AbPB, AbDM) and CONDITION (Pre-task baseline, within-task baseline, Grasp Observation), for PG and WHG separately.

To assess MEP modulation during the observation of grasp, we first compared MEPs (normalized to the within-task baseline) across baseline and grasp observation conditions via three-way rmANOVA with factors of CONDITION (Baseline, Grasp Observation) × MUSCLE (1DI, AbPB, AbDM) × GRASP (PG, WHG)). To determine whether observation of grasp modulated MEPs in a grip-specific manner, we also calculated MEP ratios using the raw, unnormalized MEPs. To obtain MEP ratios, average PG MEPs were divided by the corresponding average WHG MEPs for each condition and subject separately. A resulting ratio >1 for a given condition therefore indicated relative facilitation of PG compared to WHG, ratios of <1 indicated relative facilitation of WHG compared to PG, and a ratio equal to 1 indicated no difference between the grasps. These ratios are therefore independent of any baseline measure. We compared MEP ratios (PG : WHG) within each experiment via two-way rmANOVA (CONDITION × MUSCLE).

### Direct comparison of the “Go only” and “Go & NoGo” experiments

To assess whether grasp-specific modulation was strengthened in one experiment vs, the other, we computed the between-muscle difference of Grasp Observation MEP ratios for the two experiments and compared these differences via an unbalanced 2-way ANOVA (MUSCLE-PAIR × EXPERIMENT).

### Influence of the Go only block on the CSE during Go & NoGo block

To assess whether the CSE modulation in the “Go & NoGo” experiment was influenced by the performance of the preceding “Go only” experiment in the second group of subjects, we compared CSE modulation across the two subject groups in the “Go & NoGo” experiment, for each object and muscle separately, via 2-way rmANOVA with a within-subject factor of TMS CONDITION (Grasp, NoGo), and between-subject factor of GROUP (1 or 2).

For all ANOVAs, post-hoc comparisons were conducted where deemed appropriate and corrected for multiple comparisons (Tukey’s honestly significant difference, unless otherwise stated). P-values < 0.05 were considered significant. In cases of non-sphericity for rmANOVAs, Greenhouse-Geisser corrected p-values were used.

Post-hoc power analyses were conducted using G*Power (Version 3.1).

## RESULTS

In this study, we investigated how CSE modulation during action observation is affected by task requirements to refrain from self-movement. Subjects observed grasping actions embedded within a Go-NoGo paradigm, which required them to suppress their own movement on most trials. One group of subjects also observed grasping actions in a block without any NoGo condition (“Go only” experiment), prior to completing the Go-NoGo paradigm, to determine whether the presence of the NoGo condition influenced the level of CSE modulation during action observation.

### Task behavior

Reaction times (GO-HPR) across subjects were comparable for the two grasps (461 ± 20 ms (PG), 450 ± 12 ms (WHG), mean ± standard error across subjects). Movement times (HPR-DO) were somewhat slower for PG (679 ± 38 ms) than WHG (636 ± 45 ms), likely because the final hand shaping prior to object contact required greater accuracy, and the marble object was substantially smaller than the sphere (Figure 1a.) Experimenter reaction and movement times were typically faster than subjects, due to greater familiarity with the experiment (reaction times: 400 ± 11 (PG), 387 ± 10 (WHG), movement times: 597 ± 15 (PG), 564 ± 14 (WHG), all mean ± standard error).

### Grasp-specific EMG activity during action execution

We assessed modulation of voluntary muscle activity during subject’s own performance of the actions, to confirm that the muscles in which MEPs were being recorded were differentially involved in executing the actions. During action execution, PG elicited greater activity in 1DI and AbPB muscles, whereas WHG elicited greater activity in AbDM (Fig. 1d), as shown by a grasp × muscle interaction (F(2,26) = 31.1, p = 1.46×10^−7^, ηp^2^ = 0.71) and post-hoc comparisons of the two grasps within each muscle (1DI: p = 0.057, AbPB: p = 0.039, AbDM: p = 1.28×10^−9^). Pairwise comparisons within grasp showed that 1DI and AbPB EMG activity was not different (p = 0.99), but both muscles had significantly greater PG activity, and significantly lower WHG activity, respectively, than AbDM (all p < 0.0001).

### Non-specific facilitation relative to pre-task baseline

We compared the amplitude of MEPs evoked during the pre-task baseline to those evoked in the within-task baseline and Grasp Observation conditions during the observation task, across all 14 subjects. For both grasps, this revealed a significant main effect of condition (PG: F(2,26) = 19.15, p < 0.0001, ηp^2^ = 0.60, WHG: F(2,26) = 15.87, p < 0.0001, ηp^2^ = 0.55). The main effect of muscle and interaction effects were not significant for either grasp (all p > 0.05). MEP amplitudes were significantly smaller during the pre-task baseline (Fig. 2) compared to both the within-task baseline (PG: p = 0.00038, WHG: p = 0.00056), and Grasp Observation (PG: p = 0.0019, WHG: p = 0.016). In addition, there was weak suppression during Grasp Observation relative to within-task baseline (PG: p = 0.064, WHG: p = 0.047).

**Figure 2.**
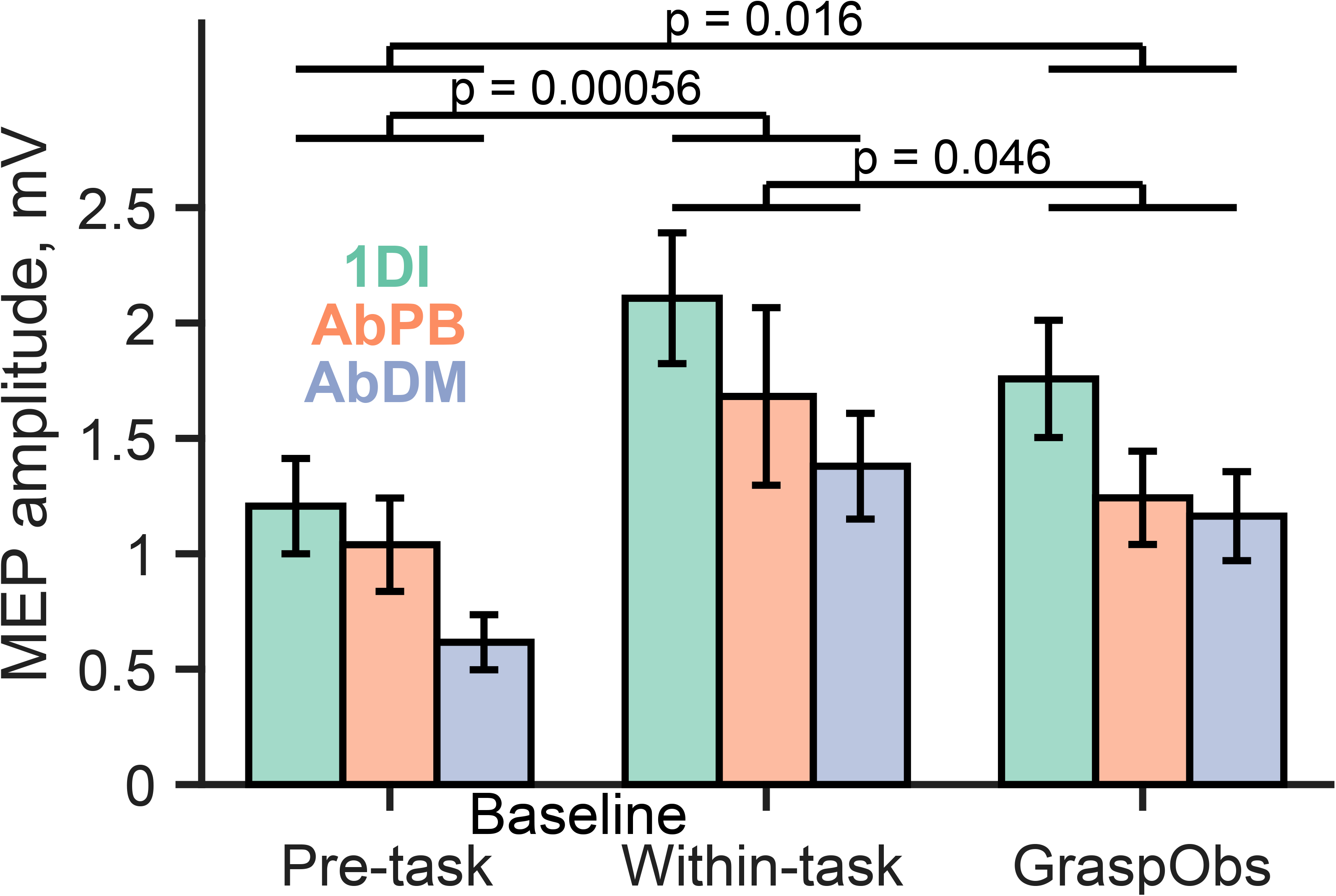
Average raw MEP amplitude across all 14 subjects for pre-task baseline, within-task baseline, and Grasp Observation (whole-hand grasp), for each muscle. Bars show mean ± SEM. P-values denote post-hoc tests between conditions.

As the comparison to the pre-task baseline predominantly exhibited broad increases in excitability, we used the within-task baseline for subsequent analyses, so that any changes relative to this baseline could be more readily attributed to the observation of action (see Discussion).

### Non-specific suppression of grasp observation MEPs in “Go & NoGo” task

To assess the modulation of MEPs during the different task conditions, we compared normalized MEPs during the “Go & NoGo” task (Fig. 3a–c) across baseline and grasp observation conditions, grasps (PG and WHG), and muscles, via 3-way rmANOVA. Although we found no significant effects with all subjects grouped together (all main effects and interactions p > 0.10), the first group of subjects showed a trend of suppression during grasp observation relative to baseline (main effect of TMS condition (F(1,6) = 5.27, p = 0.061, ηp^2^ = 0.47)). Post-hoc tests within each muscle showed that the grasp condition was suppressed relative to baseline in AbPb (mean difference −0.29, p = 0.04) and AbDM (mean difference −0.21, p = 0.02), with a weaker suppression in 1DI (mean difference −0.09, p = 0.40). A 2-way rmANOVA (condition × muscle) on the ratio of PG : WHG MEPs (Fig. 3d) showed no significant effects (all main effects and interactions p > 0.6), indicating that suppression was not grasp specific.

**Figure 3.**
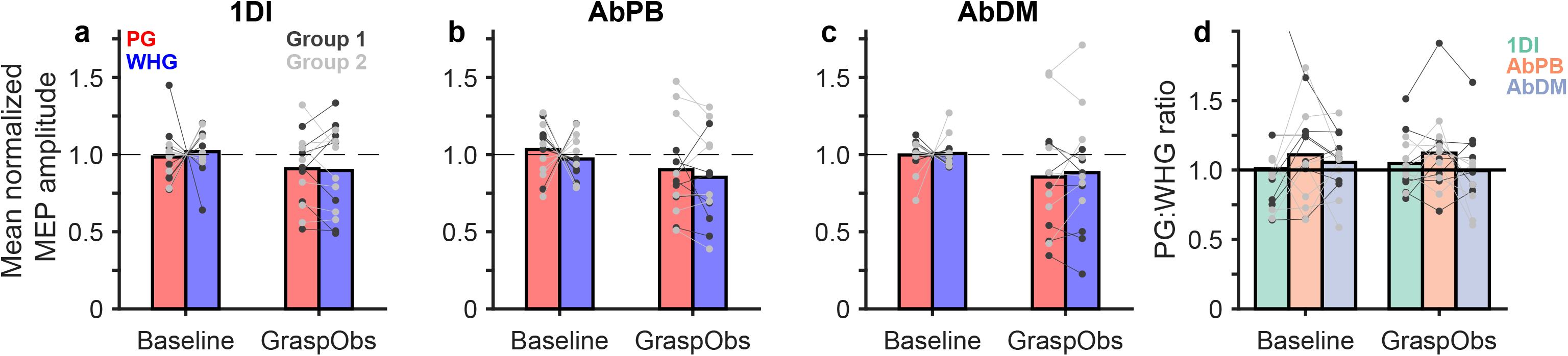
“Go & NoGo” block (n=14). **a-c.** Average normalized MEP amplitude for Baseline and Grasp Observation conditions for each object and muscle. **d.** Ratio of precision grip to whole-hand grasp (PG : WHG) MEPs. Bars show the mean, and individual subjects are shown by grey lines (dark grey: Group 1, light grey: Group 2).

### Grasp-specific modulation of MEPs in Go only experiment

We hypothesized that the presence of the NoGo condition may have increased the emphasis of the task on suppression of one’s own actions instead of observation of other’s actions (see Discussion), thus diminishing grasp observation-related MEP facilitation, or even producing weak suppression relative to baseline (Fig. 2 and Fig. 3). In the seven subjects tested later, we therefore examined this possibility by measuring CSE changes during a block without the NoGo condition, which was always performed before the “Go & NoGo” block. In addition to the absence of the NoGo condition (which would emphasise movement suppression in the absence of any observation), this block also included an increased proportion of execution trials.

In this “Go-only” experiment (Fig. 4a–c), there was a 3-way interaction between TMS condition, object, and muscle (F(2,12) = 4.53, p = 0.034, ηp^2^ = 0.43) in analysis of the MEPs normalized to the within-task baseline. In the analysis of MEP ratios, designed to better distinguish grasp-specific effects^23^, there was a significant TMS condition × muscle interaction (Fig. 4d, F(2,12) = 4.74, p = 0.030, ηp^2^ = 0.44).

**Figure 4.**
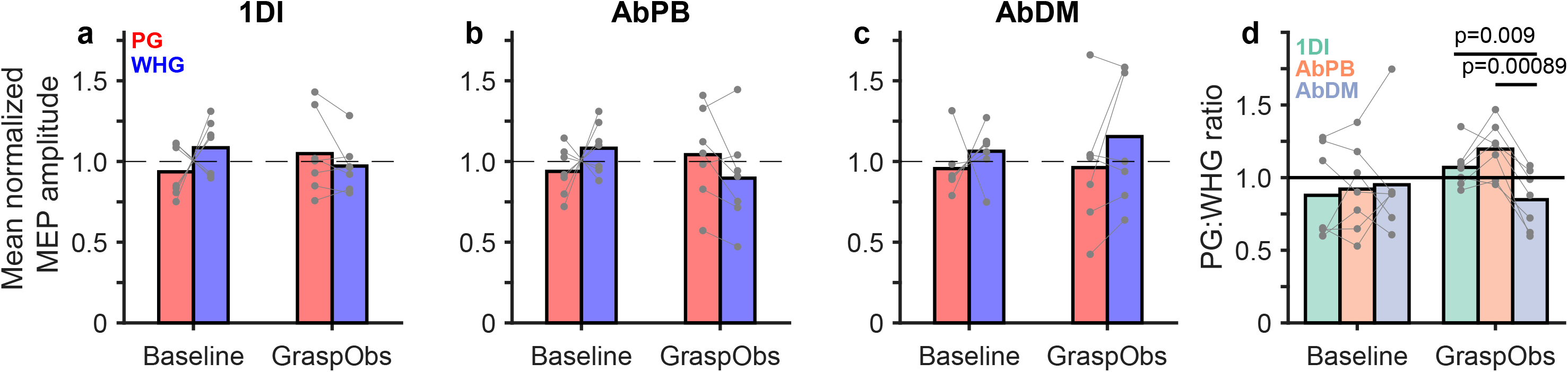
“Go only” block (n=7). **a-c.** Average normalized MEP amplitude for Baseline and Grasp Observation conditions for each object and muscle. **d.** Ratio of precision grip : whole-hand grasp (PG : WHG) MEPs. All colour conventions as in Figure 3. Bars show the mean, and individual subjects are shown by grey lines. Corrected p-values shown from statistically significant post-hoc tests following 2-way ANOVA on PG : WHG ratios.

There was no significant difference in MEP ratios between muscles in the Baseline condition (all pairwise comparisons p > 0.9). During Grasp, 1DI and AbPB ratios were not different from each other (p = 0.31), but both were significant larger than AbDM (p = 0.0090 and p = 0.00089 respectively), indicating that PG was more strongly facilitated in these two muscles, while WHG MEPs were facilitated in AbDM.

### Direct comparison of the Go only and Go & NoGo experiments

The “Go only” experiment showed a trend for increased MEP ratios in 1DI and AbPB and a decreased ratio in AbDM compared to the “Go and NoGo” experiment. To determine whether this divergence across the muscles was strengthened in the “Go only” experiment, we computed the difference in the ratios across each pair of muscles and compared these values (Fig. 5). We found main effects of experiment (F(1,57) = 4.70, p = 0.034, ηp^2^ = 0.08) and muscle pair (F(2,57) = 10.31, p = 1.5×10^−4^, ηp^2^ = 0.27), with no significant interaction (F(2,57) = 3.16, p = 0.0502, ηp^2^ = 0.10). Bonferroni-corrected post-hoc tests revealed that the AbPB-AbDM ratio difference was significantly greater in the “Go only” experiment than in “Go-NoGo” (mean increase = 0.256, p = 0.033). 1DI-AbDM and 1DI-AbPB differences were not significantly different across the two experiments (both p > 0.05). Independent t-tests confirmed that the 1DI-AbDM and AbPB-AbDM ratio differences were different from zero only in the “Go only” experiment (t(6) = 4.58, p = 0.0226 and t(6) = 7.19, p = 0.0022 following Bonferroni correction, respectively).

**Figure 5.**
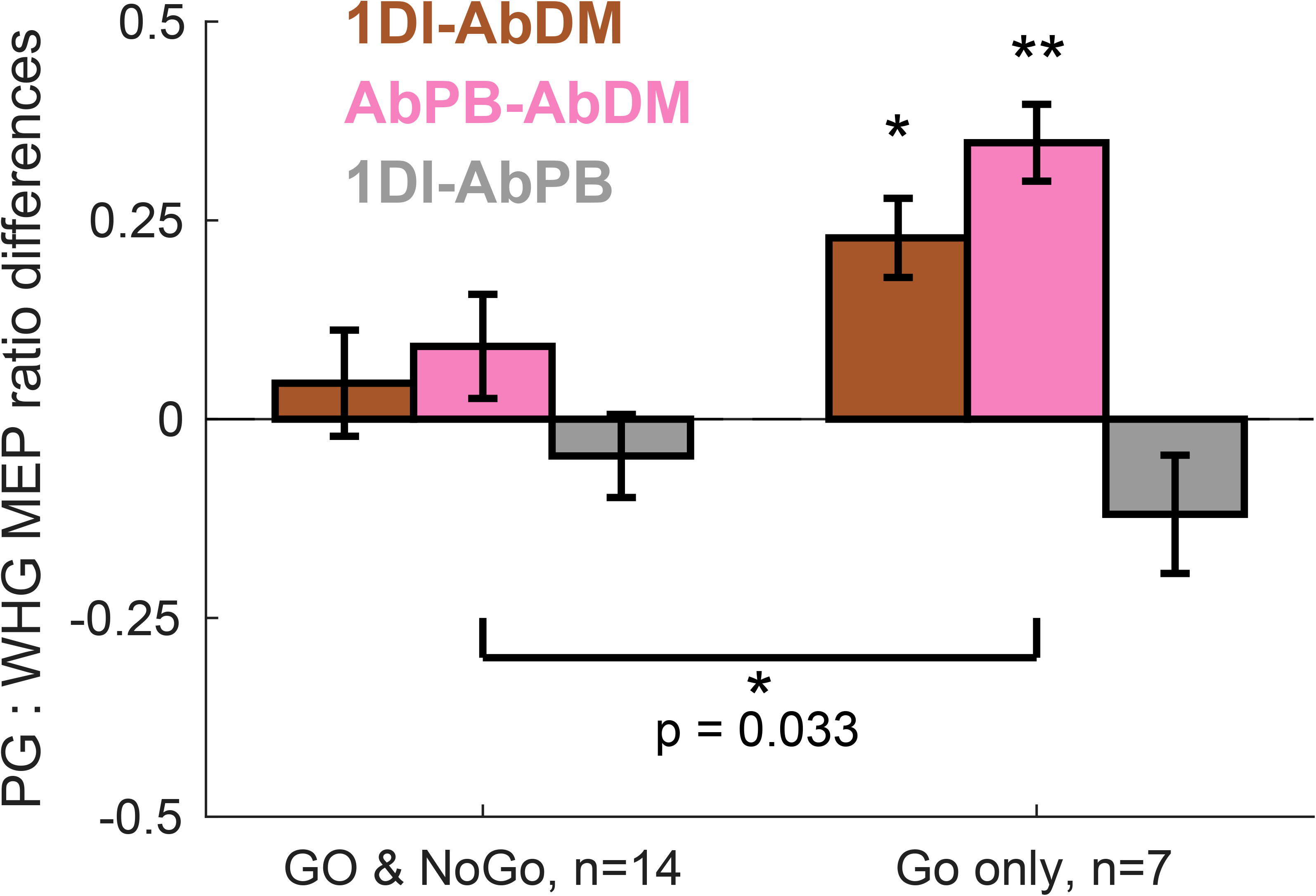
Differences between three muscle pairs (1DI-AbDM (brown), AbPB-AbDM (magenta), 1DI-AbPB (grey)) of Grasp Observation MEP ratios ((PG : WHG)), shown for “Go & NoGo” experiment (n=14), and “Go only” experiment (n=7). Data presented as mean±SEM. AbPB-AbDM difference was significantly greater in “Go only” experiment (p = 0.033). 1DI-AbDM and AbPB-AbDM differences were significantly different from zero in “Go only” experiment (two-tailed t-test * p < 0.05, ** p < 0.001).

### Influence of the Go only block on CSE modulation during Go & NoGo block

It is possible that the initial “Go-only” block performed by the second group of subjects could have influenced modulation in the subsequent “Go & NoGo” task. Since we also recorded MEPs in the NoGo condition, we compared MEPs in the Grasp observation and NoGo conditions in the “Go & NoGo” experiment across the two groups of subjects (Fig. 6) – those who performed only the “Go & NoGo” experiment (Group 1, dark grey), and those who performed the “Go only” experiment beforehand (Group 2, light grey). Figure 6 demonstrates a general pattern (for the grasp in which each muscle is preferentially engaged during execution (Fig 1d.)) in which Grasp Observation MEPs tend to be less suppressed than NoGo MEPs in Group 2, but more suppressed in Group 1. In the 1DI muscle, preferentially active during execution of PG, there was a significant TMS condition × group interaction for PG (F(1,12) = 5.10, p = 0.043, ηp^2^ = 0.11), and PG Grasp Observation MEPs were >20% greater than NoGo MEPs in Group 2 (p = 0.023). Group 1 showed the opposite pattern, although the difference between Grasp Observation and NoGo MEPs was not significant (p = 0.57). The AbDM muscle, preferentially active during execution of WHG, showed a significant TMS condition × group interaction for WHG (F(1,12) = 6.12, p = 0.029, ηp^2^ = 0.25), but not PG (F(1,12) = 1.00, p = 0.34, ηp^2^ = 0.05). In post-hoc tests, the differences between the Grasp Observation and NoGo condition did not reach significance (Group 1: p = 0.074, Group 2: p = 0.15). In AbPB, interaction effects for both objects were not significant (F(1,12) = 3.48, p = 0.087, ηp^2^ = 0.09, WHG: F(1,12) = 2.81, p = 0.12, ηp^2^ = 0.06), although a similar shift in suppression from the Grasp Observation to the NoGo condition was apparent in Group 2.

**Figure 6.**
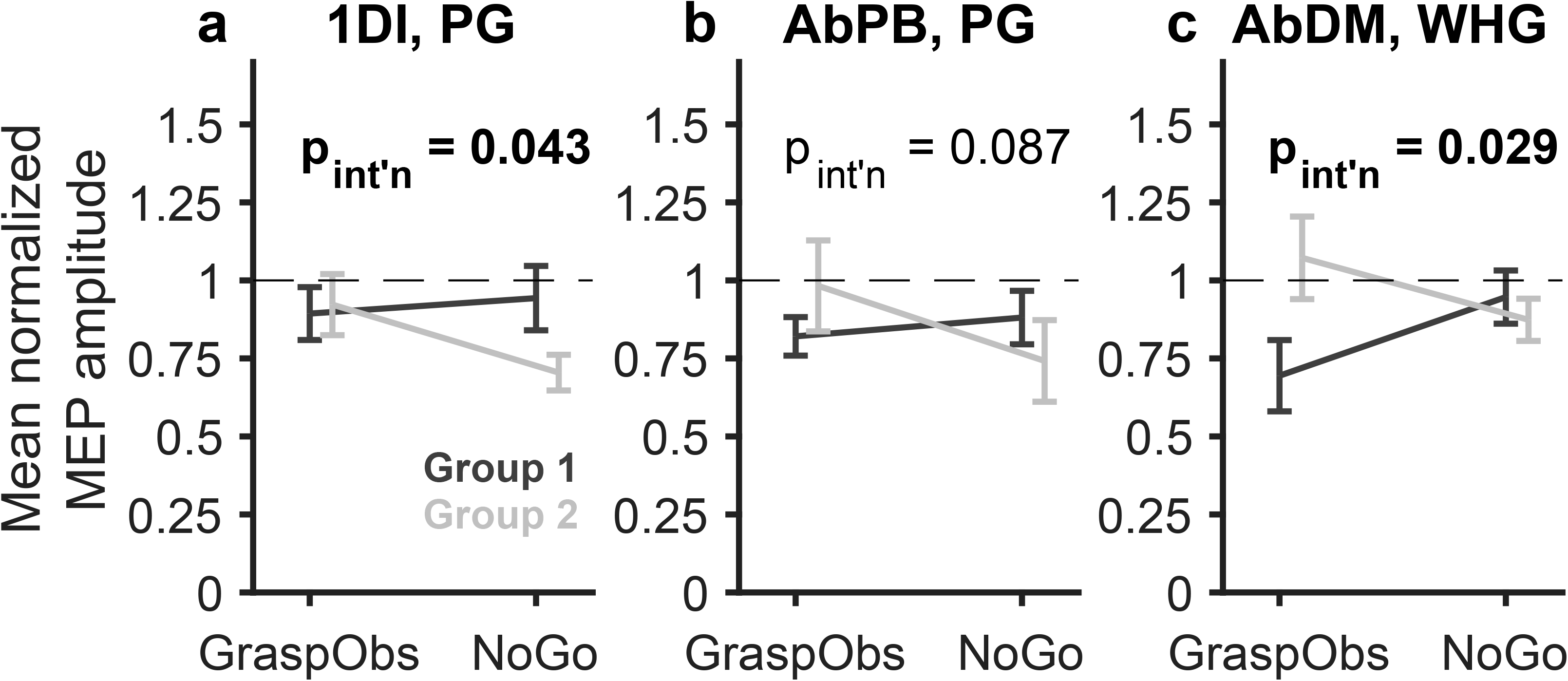
Grasp Observation and NoGo condition normalized MEPs during “Go & NoGo” block for each group of subjects (dark grey: group 1, light grey: group 2). Group 2 points are offset slightly along x-axis for clarity. For each muscle, MEPs are shown for the “preferred” grasp i.e., the one which produced greater EMG activation during action execution (Fig. 1c,d; PG for 1DI and AbPB muscles, WHG for AbDM). Data presented as mean ± SEM, and p-values are shown for interaction of condition and group (* p < 0.05, n.s. not significant)

### Post-hoc power analyses

To estimate the probability that our sample sizes enabled us to detect genuine interaction effects in our rmANOVAs, we conducted post-hoc power analyses using G*Power 3.1 ^24^. In all cases, we set α = 0.05, and non-sphericity correction ε = 0.75 for >2 measurements (otherwise ε = 1). Using a fully within-subject design (one group) with three measurements (two degrees of freedom), and correlations between repeated measures of 0 to 0.5, a strong effect size (partial eta squared ηp^2^ = 0.4, comparable to values obtained in several analyses), and n = 14 subjects (as used in analyses in Figures 1d, 2, and 3) provided an estimated achieved power of 0.98 - 0.99. For n = 7 subjects (as used in Figure 4), the same values yielded an estimated power of 0.74 – 0.96. For a relatively weak effect size (ηp^2^ = 0.05), estimated power was 0.19 – 0.35 for n = 14 subjects, and 0.11 – 0.17 for n = 7 subjects. For the within-subject rmANOVAs, this indicates that our sample size was likely sufficiently powered to detect relatively large effect sizes where they existed, but unlikely to detect weak effects. Using a within-between design with 2 groups and 2 measurements, n =14 subjects (as used in Fig. 6) and a correlation between repeated measures of 0.5, the estimated achieved power was 0.63 for ηp^2^ = 0.1, and 0.82 for ηp^2^ = 0.15.

## DISCUSSION

In this experiment, we aimed to assess the extent to which corticospinal excitability (CSE) during the observation of naturalistic grasping actions is modulated by the requirement of neuronal circuits involved in suppression of one’s own actions. Specifically, when action observation was embedded within a Go-NoGo paradigm, we hypothesized that grasp-specific modulation during action observation might be attenuated or abolished, since the emphasis of the task increasingly shifted towards global suppression of self-movement.

### Action execution recruits muscles in a grasp-specific manner

An important consideration when ascribing modulation during action observation to the mirror neuron system is that execution of the same actions produces muscle-specific activity, which shows some degree of congruence with changes during action observation^7,9^. This was a key aspect of the study by Fadiga and colleagues^7^, which reported facilitation of MEPs in three of four muscles (1DI and two arm muscles, but not a thumb muscle) during observation of arm movements, and recruitment of these same three muscles during action execution.

Despite this, a number of studies using TMS to probe CSE during the observation of naturalistic actions do not always include an ‘action execution’ condition^9^. The complex activation patterns of hand and arm muscles during different reach to grasp actions^25,26^, mean that, for example, it should not simply be assumed that AbDM is more involved in WHG than thumb and index finger muscles^9^.

In the grasping task used in the present study, PG and WHG execution reliably produce different levels of activation in the muscles we recorded (Fig. 1c,d). The 1DI and AbPB muscles showed greater activity during PG, which is consistent with their role in opposition of thumb and forefinger, whereas AbDM, a little finger abductor, showed much higher activity during WHG. This is in line with previous comparisons of EMG activity during PG and WHG performance, as well as in the excitability of PMv-M1 interactions and cortico-cortical inputs to M1 during preparation of different grasps^27^.

### Non-specific increase in CSE relative to pre-task baseline

In our task, ‘baseline’ CSE was probed while subjects were at rest in between task blocks, and also at the beginning of some trials, when subjects were still at rest and not observing any grasping action but engaged in the task. Our finding of a general (up to twofold) increase in CSE during the task compared to the pre-task baseline (Fig. 2) is quantitatively consistent with other recent results, which have shown significant increases in task MEP amplitude relative to a pre-task baseline, and much smaller changes relative to a within-task baseline^18,20^.

This non-specific increase may have been a consequence of increased attention or arousal during the task period. Although the relationship between gaze and the activity of mirror neurons is variable^28^, gaze is often used as a proxy for determining where subject attention is directed. One recent study found that direction of gaze towards the target object during observation was a significant predictor of increased MEP amplitude^29^. Similarly, D’Innocenzo and colleagues reported that fixation of gaze on a point where maximal detection of biological motion would occur produced increased MEP amplitudes relative to a free gaze condition^30^. Appropriate baseline selection is essential for determining the direction of CSE modulation during action observation, and may partly explain some of the conflicting reports of facilitation or suppression during action observation^9^. In light of the above, and because it is more closely related to the baseline typically used in assessment of mirror neuron firing rates^3^, we considered the within-task baseline a more appropriate choice for quantifying the direction of CSE modulation during grasp observation.

### Grasp-specific modulation of CSE

When subjects merely executed or observed grasping actions, we found a grasp-specific modulation of excitability during observation of PG and WHG (Fig. 4), which was not apparent when the task was completed with a NoGo condition (Fig. 3). MEPs were larger in PG compared to WHG in the 1DI and AbPB muscles, but smaller in AbDM, consistent with several TMS studies which have demonstrated muscle or grasp-specific changes in excitability during action observation^7,18,31,32^. These changes are distinct from weaker CSE changes during object presentation^7,31^, driven by canonical neuron responses in premotor area F5^33^. The study by Bunday et al. found greater excitability in the 1DI muscle during observation of videos of PG compared to WHG. Notably, this modulation largely occurred while MEPs were suppressed relative to the inter-trial interval baseline. Additionally, this interaction, when controlling for the size of the observed hand, was present when viewing the hand only, but not a whole-person actor^18^. In our study, although the whole actor was visible to the subject, the LCD screen effectively ensured that the subject’s field of view around the objects when looking through the screen was limited to include only the forearm and hand of the experimenter. Our study also differed in the fact that actions were performed by a live actor, rather than viewed on video, occurred in operational and metric peri-personal space, and from a frontal, rather than side viewpoint. In macaque mirror neurons, live actions tend to produce greater activation than video-ed ones^34^, and some mirror neurons fire in a space- ^21,22^ and view-point^34^ dependent manner. Although these distinctions render precise quantitative comparisons challenging, we did find a similar grasp-specific modulation in intrinsic hand muscles during observation of PG and WHG, suggesting that under certain conditions, observation of action induces muscle-specific changes in CSE. Importantly, these changes also parallel the recruitment of muscles during performance of those same actions (Fig. 1c,d), indicating that CSE modulation during action observation reflects activation of the human mirror neuron system.

### Task context modulates CSE

In our study, we found that the presence of the NoGo condition, which required subjects to refrain from self-movement in the same way as during observation trials, attenuated grasp-specific modulation during action observation. Subjects performing the task without a NoGo condition demonstrated grasp-specific modulation of MEPs, as revealed by PG : WHG MEP ratios (Fig. 4), and this modulation was stronger than that seen in blocks performed with a NoGo condition (Fig. 5). Differences between the two groups of subjects in the Go-NoGo experiment (Fig. 6), suggest that the factors modulating Grasp observation MEPs in Go-only block may have carried forward into the subsequent Go & NoGo block, such that suppression was largely limited to the NoGo condition for the second group of subjects, rather than appearing in the Grasp Observation as it did in the first group. This between-subject effect was relatively weak, and the condition driving the interaction effect shifted across muscles. We considered the NoGo condition as providing an emphasis on movement suppression but acknowledge that the block design presents a limitation of our study - as subjects are still required to withhold movement on observation trials in all blocks, the contextual emphasis on movement suppression was not completely removed.

Taken together, these results suggest that the relative emphasis of the task on observation of action versus suppression of self-movement, determined by task instructions and an increased proportion of trials requiring movement suppression may contribute to the balance of facilitation and suppression seen in CSE during the action observation task.

On the face of it, the rapid, automatic onset of activity in mirror neurons during action observation in a circuitry extending from parietal to frontal areas implies that bottom-up visual input is the predominant driver of mirror neuron activity. However, there is a wealth of evidence that context cues play an equally, if not more, important role. In macaque mirror neurons with ventral premotor cortex, knowledge of a grasping action taking place, even if is not visible to the observing monkey, is sufficient to elicit activity in many mirror neurons^35^. Similarly, modulation of CSE in humans can persist even in the absence of direct visual input^13,36^. On the other hand, unexpected kinematics can abolish CSE changes during action observation^8^, and although kinematic cues are sufficient for CSE modulation according to weight during observed object lifting^16,37^, incongruent cues (e.g. heavy weights labelled as light) also abolish MEP facilitation, and can even induce overall suppression^16^. Alternatively, incongruent weight labels replace the weight-based modulation with a general ‘expectation-monitoring’ one^17^. This study further showed that virtual lesioning of posterior superior temporal sulcus can restore weight-based modulation, whereas lesion of dorsolateral prefrontal cortex abolished both forms of modulation, supporting the idea that CSE modulation is susceptible to higher-level cognitive influence^17^.

Another recent study comparably found that reliable contextual cues were sufficient to elicit muscle-specific CSE changes when visual kinematic cues were absent, and that CSE modulation was strongest when both contextual and kinematic cues were present^19^. In the authors’ framework, M1 is the site of convergence for bottom-up kinematic cues regarding the actual observed action, presumably arriving via the parieto-frontal grasping circuitry which houses mirror neurons^38^, and top-down contextual cues or priors. These two streams of information may be combined within a Bayesian predictive coding framework^39^, enabling the observer to infer, or predict, the outcome or intention of the observed movement^8^.

The interactions between movement suppression during action observation and during NoGo in the present task appear to contrast with a previous study of action observation and NoGo responses in macaque mirror neurons^40^ where ‘execution NoGo’-responding neurons and mirror neurons rarely overlapped. However, these neurons were recorded in ventral premotor area F5, where population-level mirror activity is distinct from that in M1^10^. Although “passive” trials (Observation/NoGo) in the present task were more frequent than in a conventional Go-NoGo task for technical reasons (to be able to collect MEPs in a reasonable time frame), subjects were encouraged to prepare action through the interleaved trial design. In addition, information about trial type (execution, observation, NoGo) was provided at the equivalent time moment on each trial, such that subjects could not predict the trial type before this timepoint, whereas the mirror neuron study examined execution and observation responses in separate blocks. The presence of the two objects in peri-personal space may also have engendered a closer relationship between action observation and NoGo, as the possibility to interact with the objects increases the relevance of appropriate and timely movement suppression. In line with this, some mirror neurons are known to preferentially encode peri-personal space^21^, including in an operational manner, suggesting that the possibility of object interaction is a salient contextual factor contributing to modulation during action observation.

Our results indicate that although a grasp-specific ‘motor resonance’ effect can be elicited by action observation, this effect is not robust, and its magnitude and direction are influenced by task instructions to suppress action. This sits within a growing body of evidence indicating that the overall modulation of CSE during action observation is the net result of the interplay between bottom-up visual input and top-down cognitive contextual cues and priors. Future studies with a larger subject cohort and within-block designs are needed to disentangle the effects of explicit and implicit contextual requirements related to movement withholding on the modulation of CSE during action observation.

## Acknowledgements

The authors thank Adam Keeler and Julie Savidan for assistance with data collection and Spencer Neal, Jonathon Henton, Chris Seers, and for technical assistance.

## Notes

**Conflict of Interest: ‘**Authors report no conflict of interest’

**Funding sources:** S.J.J was funded by a Brain Research UK Graduate Student Fellowship. A.K. was funded by the Wellcome Trust, grant number 102849/Z/13/Z and BBSRC grant BB/P006027/1.

### Competing Interest Statement

The authors have declared no competing interest.

### Summary of Updates

Text rewritten after reviewers comments

